# A mechanistic basis for CD8+ T cell expansion sensitivity as a predictor of HIV post-treatment control

**DOI:** 10.64898/2026.08.28.747758

**Authors:** Tin Phan, Nicole Pagane, Jasmine Kreig, Aurelien Marc, Macauley Locke, Michael J. Peluso, Demi A. Sandel, Amelia N. Deitchman, Rachel L. Rutishauser, Steven G. Deeks, Ruian Ke, Ruy M. Ribeiro, Alan S. Perelson

## Abstract

A key goal in HIV-1 cure research is to understand why some individuals control viral rebound after stopping antiretroviral therapy (ART). Recent human studies have identified responding CD8+ T cells expressing Ki-67 and the transcription factor TCF-1 as correlates of post-treatment control, but the mechanistic basis of this association remains unclear. Using the theoretical framework of Conway and Perelson, we fit mechanistic within-host models to viral load and CD8+ T cell data from 9 individuals in a combination immunotherapy trial following ART interruption. Although Ki-67 and TCF-1 measurements were not used for fitting, the inferred effector cell expansion sensitivity, i.e., the responsiveness of effector expansion to low antigen levels, shows a strong linear relationship with Ki-67 and TCF-1 levels at rebound (Pearson’s r ≈ 0.8). Building on this, we show analytically that the post-rebound viral load set point is inversely proportional to the effector cell expansion sensitivity, and thus strongly correlates with cycling (Ki-67+) CD8+ T cells (r ≈ –0.8) at rebound, and a subset that expresses TCF-1 (r ≈ –0.9). In effect, individuals with a larger proportion of CD8+ T cells responding to viral rebound, and a greater representation of TCF-1 expressing cells within the responding subset, achieve markedly lower viral set points through a higher effector cell expansion sensitivity. This mechanism is consistent with prior modeling in a non-intervention ATI setting, suggesting it may generalize across more rebound contexts. Our results provide a mechanistic explanation why both Ki-67+ responding CD8+ T cells and their TCF-1-expressing subset predict post-treatment control, linking clinical correlation to its underlying cause and highlighting Ki-67 and TCF-1 as potential early biomarkers of HIV immunotherapy success.

## I. Introduction

A major goal in HIV cure research is the suppression of virus after antiretroviral therapy (ART) interruption. This is in part motivated by individuals, known as post-treatment controllers (PTCs), who stay in remission for months to years following analytical ART interruption (ATI), in contrast to the rapid viral rebound of non-controllers (NCs). The mechanistic difference between PTCs and NCs is not fully understood; nevertheless, several analyses have pointed to differences in the immune response [1–7] and characteristics of the latent reservoir [8–11], as primary drivers of extended viral control. Several groups have experimented with ways to enhance the likelihood of viral control post-ATI by means of immune-modulators [4,6,12], broadly neutralizing antibodies (bNAbs) [13–15], and latency reversing agents (LRA) [16–20].

In a recent single-arm clinical trial (the UCSF-amfAR combination immunotherapy study [NCT04357821]), Peluso, Sandel and Deitchman et al. [4] administered multiple immunotherapies (an HIV Gag conserved element vaccine regimen, a TLR-9 agonist, and bNAbs) over a 34 week period and demonstrated viral control in 7 out of 10 people with HIV (PWH) – a substantially higher success rate than historically seen in prior ATI studies [2,13,14,21–25]. One participant remained aviremic for 18 months post-ATI and was deemed a post-treatment controller before resuming ART. The other 6 exhibited viral rebound and then maintained a low viral set point and were termed post-intervention controllers (PICs). They found that an early, robust expansion of activated CD8+ T cells in response to rebounding virus was associated with viral control after ATI, alluding to the importance of CD8+ response in post-intervention control. In particular, the frequency of non-naïve CD8+ T cells in cell cycle (Ki-67+) that co-expressed the T cell memory transcription factor T cell factor (TCF)-1 measured at the first peripheral blood mononuclear cell (PBMC) sampling time point after viral rebound was significantly higher in PICs than in NCs [4]. Notably, despite all individuals receiving the same sequence of immunotherapy interventions, viral rebound dynamics varied widely, ranging from rapid rebound and high set points to prolonged low-level (<1000 copies/mL) viremia. This suggests a heterogeneity in immune response effectiveness. Recent work further supports this finding, highlighting polyfunctional responses from HIV-specific stem-like CD8 T cells as a marker of post-treatment control [3,6,11,12,26,27]. However, the mechanistic link between these immune correlations and sustained viral control is not fully understood.

Mathematical models have continually been developed alongside advances in clinical HIV studies to leverage new experimental capabilities, mathematics, and computations to provide quantitative insights into the effects of treatment including immunotherapy on HIV dynamics and latency [28–36]. We previously built on the Conway and Perelson modeling framework [1] to analyze a data set from Sharaf et al. [37], where we found the expansion sensitivity of effector cells, which is a quantitative measure of their proliferative responsiveness to low antigen levels, to be a significant determinant of post-treatment control [38]. However, that study did not connect the model-inferred expansion sensitivity to any measurable immunological marker, leaving the biological interpretation of this parameter unresolved. In this study, we extend our modeling framework to quantitatively analyze the viral load and CD8+ T cell dynamics and observations by Peluso, Sandel and Deitchman et al. [4]. Our goal was to investigate the biological mechanisms underlying the observed correlations — in particular, to determine *how* TCF-1+ CD8+ T cell responses might contribute to viral control and to reconcile the heterogeneous rebound patterns within a unifying quantitative framework. Strikingly, the model-inferred expansion sensitivity, estimated solely from viral load and CD8+ T cell dynamics, correlates strongly with both the size of the responding CD8+ T cell pool (%Ki-67+) and its TCF-1+ fraction at rebound, measurements that were not used in the model fitting.

## II. Methods

### Data

The longitudinal viral load and CD8 T cell data were from the study by Peluso, Sandel and Deitchman et al. [4] that included 10 participants, each receiving combination immunotherapy with an HIV Gag conserved-element DNA vaccine plus an IL-12 prime/modified vaccinia Ankara boost, bnAbs 10-1074 and VRC07-523LS, and a TLR-9 agonist lefitolimod during ART, and a repeat bnAb infusion 48 hours before the time of ATI. After ATI, weekly measurements of viral load were collected for 24 weeks and then every other week afterward.

For model fitting, we used the longitudinal viral load and CD8+ T cell measurements post-ATI to examine the viral rebound dynamics. Hence, we did not include participant PID 60610 whose viral load was never above the lower limit of quantification (LLoQ of 30 copies/mL) in our analysis. Of the nine remaining participants, three were NCs who experienced rapid viral rebound to over 5000 RNA copies/mL [4]. The other six were PICs, who ultimately suppressed viral loads below 2000 RNA copies/mL for at least 2 months [4]. For downstream analysis, we used the following measurements at the first PBMC sampling time point post-rebound called post-R1: the %Ki-67+ cells of non-naïve CD8+ T cells (what we consider to be a proxy for the magnitude of the CD8+ T cell population “responding” to rebound [39,40]), and the %TCF-1+ cells within this responding CD8+ T cell population. This additional data was available for 8 participants and is provided in the supplemental information of reference [4]. The post-R1 sample is important because cellular measurements made at this time point reflect the state of the immune system as virus is starting to rebound, and it was collected when viral loads were similar between PICs and NCs.

## Mathematical Models

### The Conway-Perelson model

We started with the Conway-Perelson model (Eq. 1) of HIV-1 dynamics [1], a foundational framework whose variations have successfully reproduced viral load trajectories in HIV, SIV, and SHIV [38,41–43].

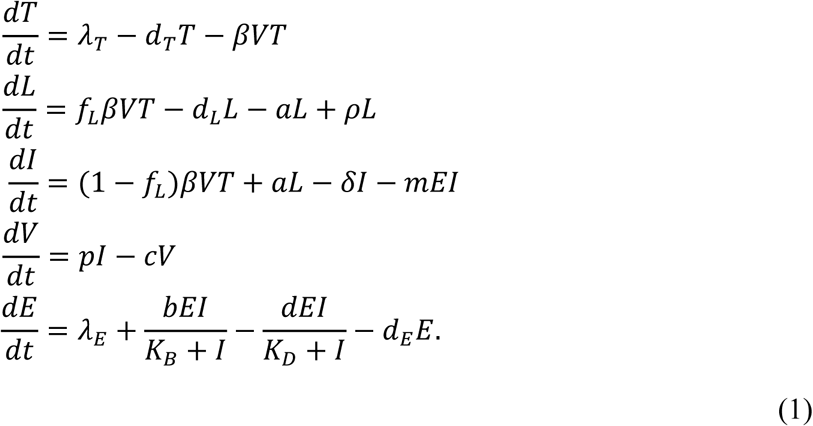

The Conway-Perelson model (Fig. 1) describes the interactions between uninfected target cells (*T*), latently infected cells (*L*), productively infected cells (*I*), virus (*V*), and effector cells (*E*). In the absence of infection, target and effector cells are maintained at their homeostatic levels by constant production at rates (*λ_T_* and *λ_E_*), and natural death at rates (*d_T_* and *d_E_*). Virus is cleared at per capita rate *c* and with rate constant *β* infects target cells, which subsequently become either latently infected cells (with probability *f_L_*) or productively infected cells (with probability 1 − *f_L_*). Latently infected cells proliferate at per capita rate *ρ*, die at per capita rate *d_L_*, and can reactivate to become productively infected cells at per capita rate *a*. Productively infected cells produce virus at rate *p* and die at per capita rate *δ*. They also stimulate the expansion of effector cells according to 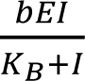, where *b* is the maximal expansion rate and *K*_B_ is the infected-cell density at which the expansion rate reaches half-maximal. We define 1/*K_B_* as the *expansion sensitivity*. Note higher sensitivity, i.e. lower *K_B_*, means effector cells are more sensitive to the level of infected cells, begin expanding at a lower antigen burden and reach their half-maximal expansion rate faster. The parameter *K_B_* is also related to the affinity of the T cells for antigen [44]. Effector cells kill infected cells with rate constant *m* and become exhausted according to 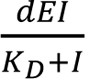, where *d* is the maximal exhaustion rate and 1/*K_D_* is the exhaustion sensitivity with *K_D_* being the infected-cell density for which the exhaustion rate is half-maximal.

**Figure 1.**
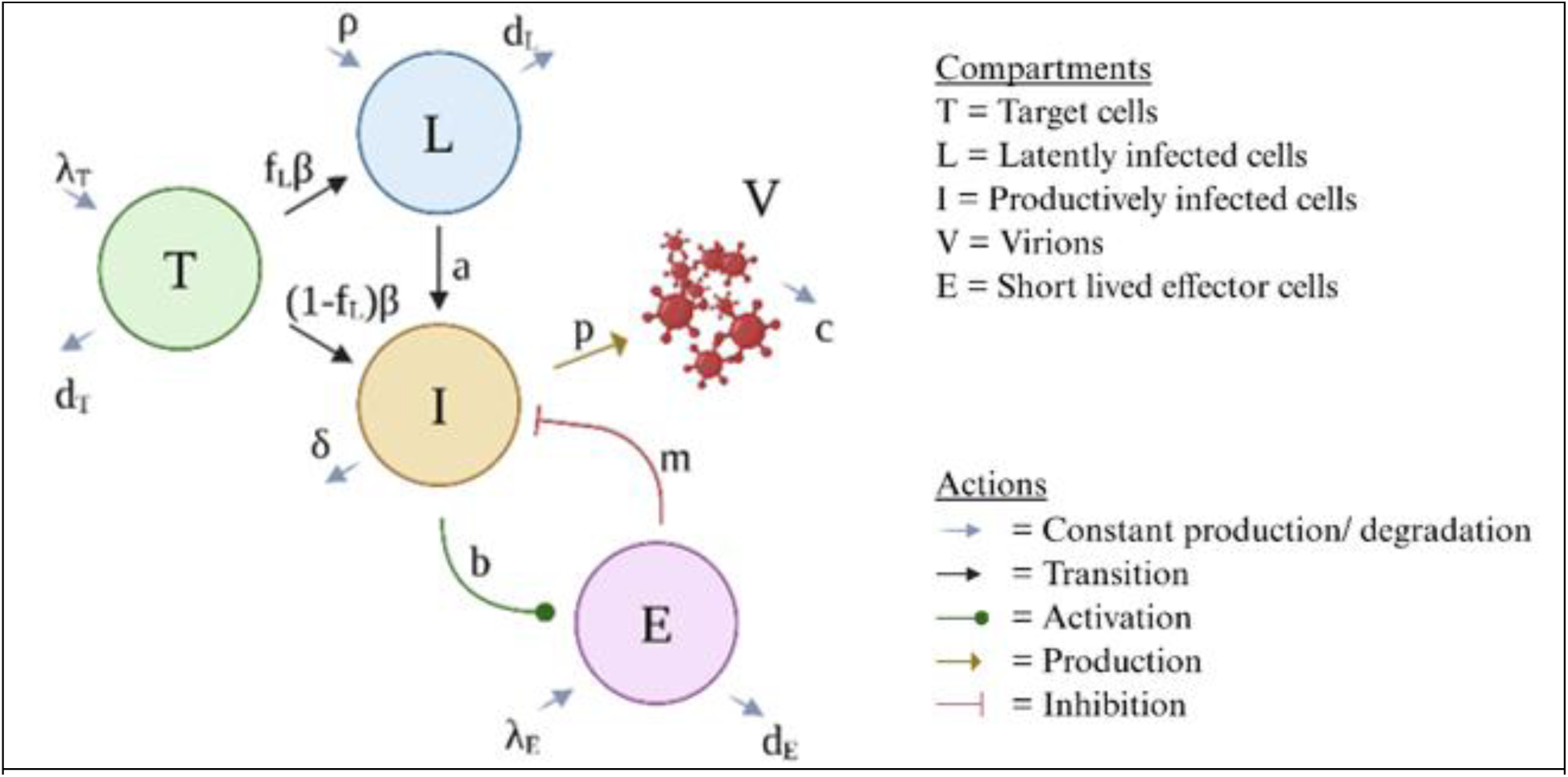
Schematic for the Conway – Perelson model.

We previously noted that the similar functional forms for the expansion and exhaustion of effector cells in the Conway-Perelson model may lead to difficulties in estimating their effects individually [38]. This is likely due to insufficient data to distinguish the various stages and effects of immune exhaustion [45]. Thus, we use a variation of the Conway-Perelson model where the exhaustion term 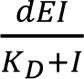 is neglected. Here, we interpret the effector cell expansion term 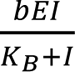 as an effective expansion term that accounts for the exhaustion effect. Note that this also alters the interpretation of *b* and *K_B_*. Instead of representing the intrinsic maximum expansion rate and half-saturation constant, respectively, they should be understood as effective expansion parameters that implicitly incorporate the dampening effect of exhaustion. The Conway-Perelson model without explicit effector exhaustion was the best performing model in our previous study [38] and is the main model studied here.

While effector cells can also mediate non-cytolytic effects that limit viral infection and production [46–50], we do not include non-cytolytic effects in the main model for a similar reason. Finally, while multiphasic declines in viral load are often observed after ART initiation in HIV, SIV, and SHIV [51–53], suggesting different infected cell death rates or mechanisms associated with different integration rates [41,54,55], we do not have the data resolution to observe these effects, so we model the productively infected cell population with a single death rate *δ*. We provide a comparison of main model variations that include these effects to justify our model choice in Table S1 Supplementary Material.

### Parameter estimation and analysis

We performed parameter estimation using a non-linear mixed effect modeling framework (software Monolix 2024, Lixoft, SA, Antony, France) to simultaneously fit the model to the viral load and CD8+ T cell data from all nine individuals. We fit the model log_10_ *V* variable to the log^10^ viral load data and the model log_10_ *E* to the log_10_ *f_E_* CD8+ T cells, where *f_E_* represents the fraction of total CD8+ T cells that the model attributes to the effector cell compartment. We do not assume that *E* is simply the circulating HIV-specific CD8+ T cells as *E* may also reflect effects of CD8+ T cells in tissues and the effects of other populations such as NK cells. In other words, we assume that the dynamics of the effector cells are reflected in the dynamics of the total circulating CD8+ T cell count over the observation window, treating the effector fraction of total CD8+ T cells as approximately constant. In our model fitting, we applied left censoring of the data for values at or below the LLoQ. Model comparison was done using the corrected Bayesian Information Criterion (BICc) [56] as reported by Monolix.

While the slow decay of the bNAbs given just prior to ATI may influence the time to rebound [4], our model accounts for these differences, forgoing modeling individual drug PK-PD to examine immune and viral factors shaping rebound dynamics once rebound is underway. First, to bypass the stochastic phase between ATI and viral rebound [1,8], we introduce a parameter *t_a_* > 0, representing the first time the virus successfully rebounds. Thus, *t_a_* absorbs participant-to-participant differences in drug washout and the stochastic establishment of successful viral rebound. Second, because the infection kinetic parameters are estimated as effective, individually fitted quantities, any residual drug effect persisting into rebound, for example, ongoing neutralization by bNAbs, is absorbed into these estimates. We assume that before time *t_a_*, the level of infectious virus and antiviral immune response were negligible. Furthermore, we assume the system is in quasi-steady state [57,58]. These assumptions lead to the following simplified initial conditions at ATI (*t* = 0): *T*(0) = *λ_T_*/*d_T_*, *L*(0) = *L*_0_, *I*(0) = 0, *E*(0) = *λ_E_*/*d_E_*. The latency reactivation rate *a* is also set to 0 before the successful rebound time *t_a_*. Once viral rebound has been initiated (at time *t_a_*), the subsequent dynamics are governed by viral replication and the immune response rather than by the reactivation rate *a*. Based on our previous study [38] where we found a covariate on *K_B_* helped the model better differentiate PTCs and NCs, we also introduce a covariate on *K_B_* to the main model here. The value and range of model parameters closely follow previous studies [1,38]. A summary of the description, range of parameter values explored, and references for all parameter values are provided in Table 1.

**Table 1.**
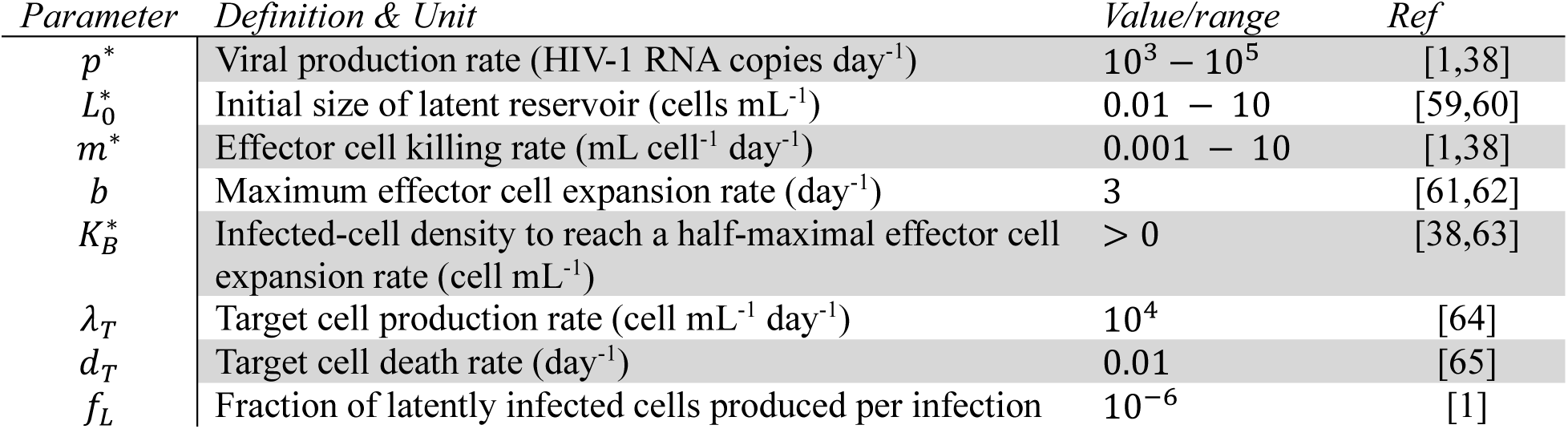
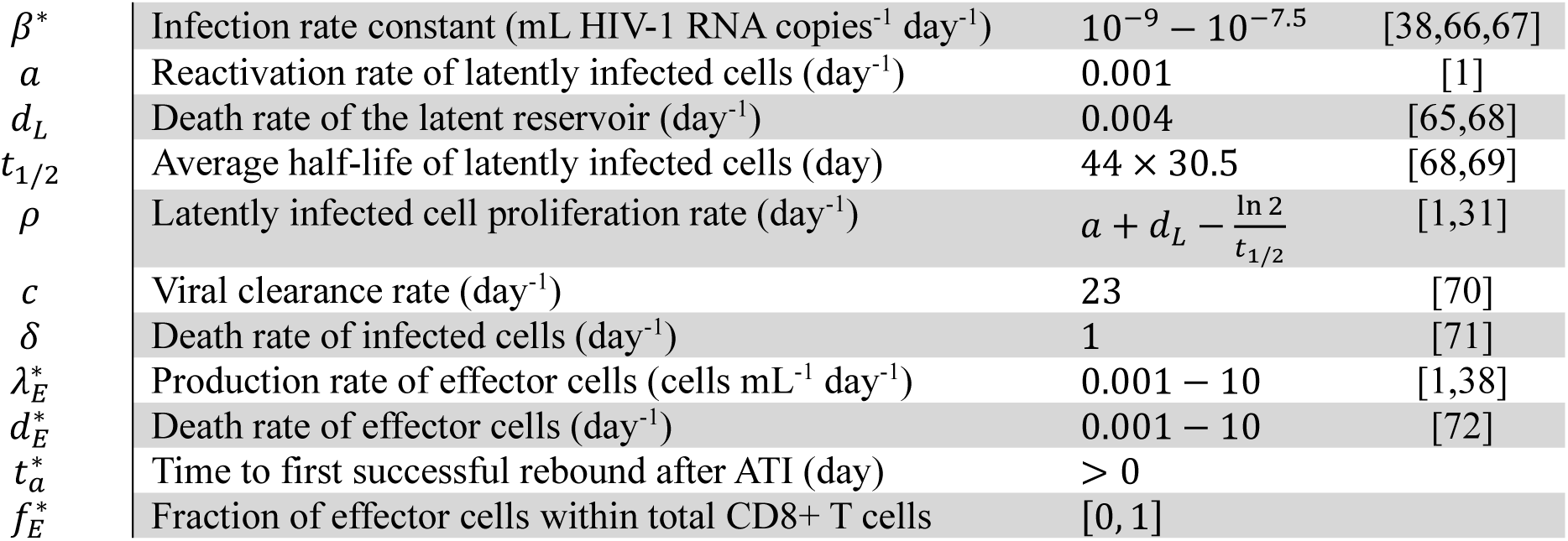
Parameter definition, range, or fixed value. The nine parameters with an asterisk are fitted using VL and CD8+ T cell data in the main model variation.

## III. Results

### Viral dynamics models recapitulate longitudinal VL data

The main model recapitulates both the heterogeneous patterns of viral rebound (Fig. 2a) and CD8+ T cells (Fig. 2b). While the pattern of rebound in NCs (PID 35933, 42894, 31092) is relatively uniform, PICs exhibit greater heterogeneity in rebound VL pattern, which is characterized by short (PID 10227, 72210) and long (PID 48758) duration oscillations, rapid settling to a viral set point (PID 47019), or a slowly increasing average VL (PID 30988). Our model captures not only overall trends but also the spectrum of controller vs. non-controller dynamics observed clinically (Fig. 2a). CD8+ T cell count data sometimes appear noisy and erratic, with instances of measurements falling completely out of trend, for example PID 47019 and 30988 (Fig. 2b). Nevertheless, the model captures the general trends well. In particular, when the dynamics are pronounced, the model fit closely follows the trend, see PID 10227, 47019, 48758, 71190, and 72210. Since the clinical definition of rebound generally depends on the time VL reaches and maintains a certain threshold [24,25,37], the model is suited to capture this characterization.

**Figure 2.**
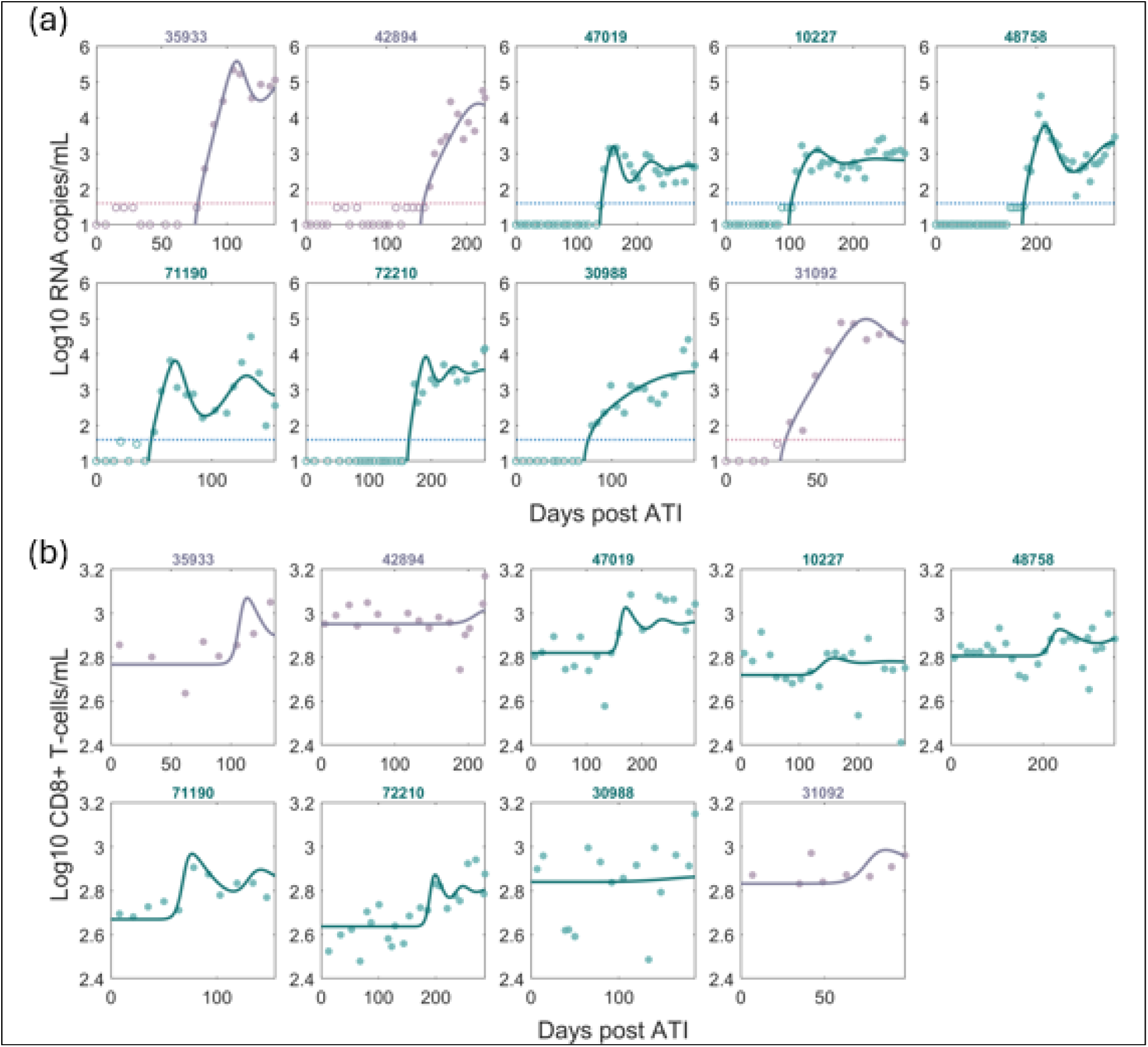
Best fit of the main model variation. Purple indicates NCs and Green indicates PICs. Filled and open circles are measurements above and below the LLoQ, respectively. Horizontal dotted lines in (a) indicate the LLoQ. (a) Model fit to VL data. (b) Model fit to CD8+ T cell data.

### Expansion sensitivity of effector cells separates PICs and NCs

Population estimates of the model parameters that were fit along with the 95% confidence interval (CI) and the range of values for individual estimates are shown in Table S2. We stratified the individual best-fit parameter values by outcome (NCs vs. PICs; Fig. 3a). Three parameters (*β*, *p*, *K_B_*) showed statistically significant differences between PICs and NCs (Mann-Whitney P<0.05). However, the median values for the infection rate *β* and the infected cell viral production rate *p* differed only marginally between groups (note the y-axis ranges). In contrast, the infected cell density at which the effector cells expand at half-maximal rate, *K_B_*, was over an order of magnitude lower in PICs than in NCs. The parameter *K_B_* alone provided a clear separation between the two groups, far exceeding the effect of any other single parameter. This supports the notion that *K_B_*, or equivalently its inverse (i.e., the effector cell expansion sensitivity) is the dominant differentiator between PICs and NCs.

**Figure 3.**
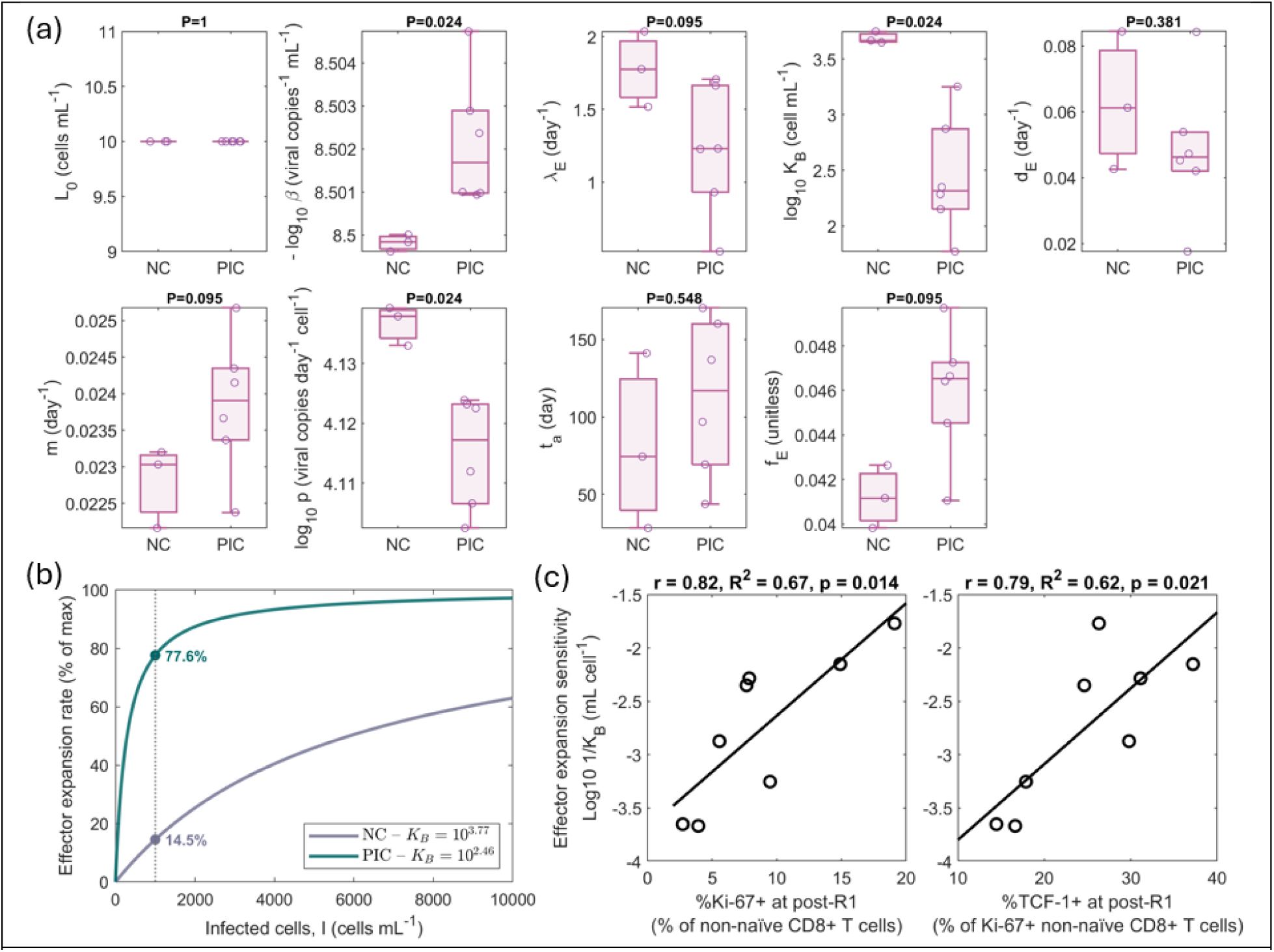
Stratification and analysis of fit parameters. (a) Individual best-fit parameters stratified by NCs (n = 3) vs PICs (n = 6) with p value calculated using the Mann-Whitney test. Individual parameter values are presented in Table S3. (b) Effector expansion rate as a function of the number of infected cells 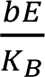. The curves for NCs and PICs were generated using their respective median values for *K_B_*, which differ by 20-fold. (c) Pearson correlation between log_10_ 1/*K_B_*and the size (%Ki-67+ of non-naïve CD8+ T cells) and stem-ness (%TCF-1+ of Ki-67+ cells) of the responding CD8+ T cell population at post-R1.

### Effector cell expansion sensitivity (1**/***K_B_*) correlates with both the size and the stem-like composition of the responding CD8+ T cell pool at the post-R1 time point

The effector cell expansion sensitivity provides a means of evaluating how rapidly effector cells expand in the presence of antigen, i.e., productively infected cells. The rate of effector cell expansion is given by 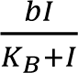, which can be rewritten as 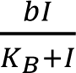, where *Z* = *I*/*K*_B_. When *Z* ≫ 1, the expansion rate is approximately *b*, the maximum expansion rate. For PICs, the mean sensitivity is 3.5 × 10^−3^, thus if *I* is ≥ 10^3^ cells/mL, the expansion rate will be greater than or equal to 3.5/4.5, or 78%, of the maximum expansion rate, whereas for NCs the corresponding expansion rate will only be at 15% of the maximum (Fig. 3b).

As shown in Fig. 3c the expansion sensitivity is also related to experimental observations not involved in our analysis. In particular, the effector cell expansion sensitivity 1/*K_B_* correlates strongly with both the %Ki-67+ of non-naïve CD8+ T cells (r = 0.82, p = 0.014) and the %TCF-1+ cells within the Ki-67+ non-naïve CD8+ T cells (r = 0.79, p = 0.021; Fig. 3c). In other words, participants whose responding CD8+ T cells (i.e., Ki-67+ during early rebound) had a more stem-like profile (i.e., a larger TCF-1+ fraction) exhibited significantly higher expansion sensitivity and ultimately better viral control. Notably, across all iterations and perturbation of the main model, where we vary the number of fitting parameters, randomize initial guesses for fitting parameters, and perturb fixed parameters, both relationships remained robust (r ≈ 0.79 for %Ki-67 vs. 1/*K_B_* and r ≈ 0.76 for %TCF-1 vs. 1/*K_B_*) (Tables S4-S5, S3 Text). These results also extended to perturbations of the model structure such as model variations with explicit effector cell exhaustion, or with non-cytolytic effector function. These findings suggest that at the post-R1 time point the size of the responding CD8+ T cell pool, indicated by %Ki67+, and its stem-like composition, indicated by %TCF-1, form a mechanistic indicator of post-ATI control in this study, in line with the idea that Ki-67 and TCF-1 mark a long-lived, highly expandable subset of antigen-specific CD8+ T cells [73–75].

We demonstrate this phenomenon in Fig. 4, where the lower values of *K_B_* (for PICs) result in earlier effector cell expansions compared to NC. Specifically, the effector cell population reaches approximately the same steady state level (log_10_ 1.5 cells/mL) about 2 weeks faster for PICs compared to NCs. Importantly, the distinction – between *sensitivity* (how quickly effectors respond to low levels of antigen) and *capacity* (the ultimate size of the effector population) – is meaningful. The steady-state effector levels differ negligibly between PICs and NCs (Fig. 4a), whereas the speed of expansion differs dramatically. This implies that the speed at which the effector population initially expands strongly influences its ability to control the infected cell burden, with the 20-fold decrease in *K_B_* for PICs vs NCs, i.e., 20-fold increase in effector cell sensitivity, corresponding to a roughly 20-fold decrease in the infected cell (Fig. 4b) and viral set points (Fig. 4c). We illustrate this point further in Fig. 4d, where faster-expanding effector cells in PICs rapidly curb the growth of infected cells, leading to a lower infected cell burden compared to NCs.

**Figure 4.**
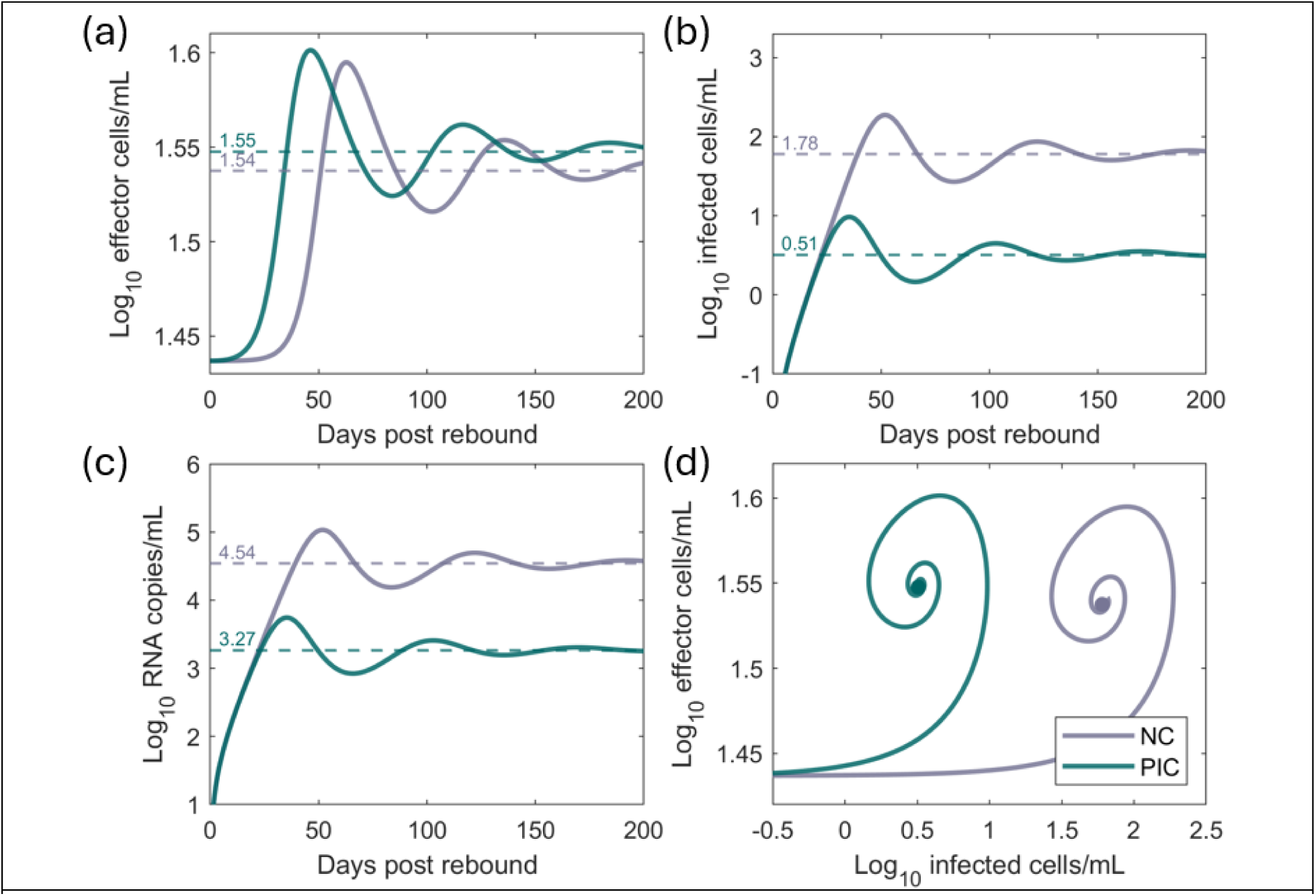
Effect of varying. *K_B_* on the effector and infected cell populations. The main model is simulated with the best-fit population parameters given in Table S2, where only *K_B_*differs between PICs and NCs. Predicted post-rebound dynamics: (a) effector cell population, *E* (b) infected cell population, *I* (c) viral load, *V*. The dashed-horizontal lines indicate the predicted steady-state levels. (d) Phase plot of effector vs. infected cells on a log scale.

### %Ki-67 and %TCF-1+ of Ki-67+ non-naïve CD8+ T cells at post-R1 correlate with the viral set point

In our previous modeling study [38], we established a simple linear relationship between *K_B_* and the viral set point post-rebound *V*_set point_ ≈ *αK_B_*, where the constant *α* is calculated based on model parameters (see S4 Text). This approximation also works for the nine individuals studied here, where we observe that the viral set points either approach or oscillate around the calculated *V*_set point_ (Fig. S1). Since 1/*K_B_* correlates with the size and stemness of the responding CD8+ T cell population (Fig. 3c), this suggests the viral set point should also negatively correlate with this measurement. To test this, we defined each participant’s viral load set point as the average viral load over the last 2 weeks of observation (about 3 measurements per participant; Fig. S2 shows the robustness to this window), rather than the more clinically oriented steady-state definition by Peluso, Sandel and Deitchman et al. [4], who averaged all viral load measurements starting 2 weeks after the rebound peak. Because our model was fit to all viral loads, its inferred set point tracks the endpoint viral load (Fig. S1). Using this set-point definition, we found strong and significant correlations between the viral set point and both %Ki-67+ at post-R1 (r = –0.79, *p* = 0.019; Fig. 5a) and %TCF-1+ fraction within those cells (r = –0.89, *p* = 0.003; Fig. 5b). The latter is substantially higher in magnitude than the –0.74 correlation reported previously [4]. This improvement in the correlation suggests that TCF-1’s impact on viral control might have been underestimated in the original analysis.

**Figure 5.**
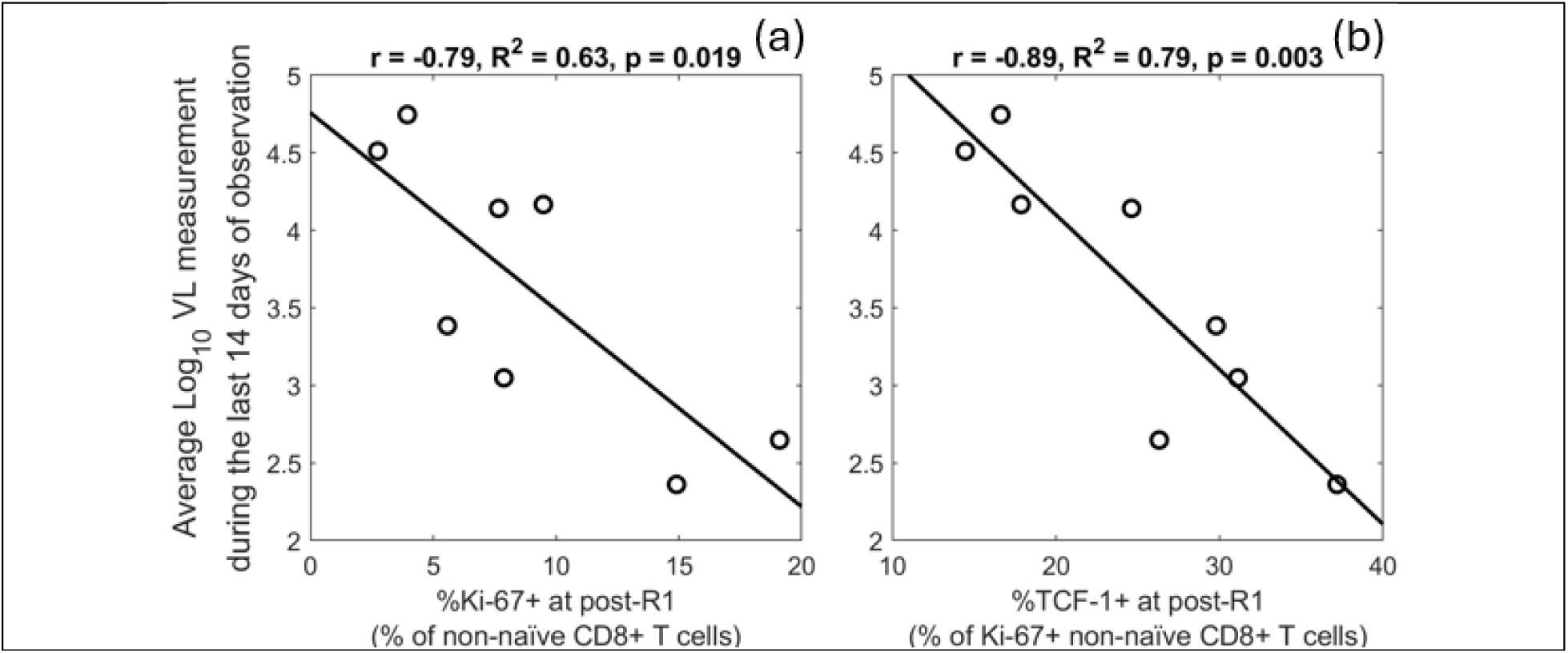
Viral set point correlations with markers of the responding CD8+ T cell population. (a) Viral set point correlation with %Ki-67+ of non-naïve CD8+ T cells at the post-R1 time point. (b) Viral set point correlation with %TCF-1+ of Ki-67+ non-naïve CD8+ T cells at post-R1. Here the viral load set point was defined as the average viral load over the last 2 weeks of observation.

## IV. Discussion

In this study, we developed dynamic models of virus-immune interactions to investigate the mechanisms behind HIV rebound control following combination immunotherapy in the UCSF-amfAR study [4]. Our analysis provides a mechanistic explanation for the clinical observations by Peluso, Sandel and Deitchman et al. [4]. We confirm that an enhanced CD8+ T cell response, specifically a high expansion sensitivity of the effector cell population to the level of HIV-infected cells, is the key determinant distinguishing post-intervention controllers from non-controllers in this combination immunotherapy trial. By accurately recapitulating the viral load and peripheral blood total CD8+ T cell count dynamics for the nine individuals in this study that had measurable viral loads after ATI, the model reveals how early immune dynamics in response to HIV rebound shape viral rebound kinetics. The model shows that individuals with greater sensitivity for effector cell expansion (as indicated by Ki-67 and TCF-1) can control rebounding HIV better, leading to a lower viral set point. This finding suggests we should reframe the concept of effector cell “expansion capacity” [75] as fundamentally a question of expansion *sensitivity*: what determines outcome is the threshold at which effectors begin responding. We show that the model-derived expansion sensitivity of effector cells correlates strongly with both the size of the responding CD8+ T cell population (Ki-67+ non-naïve CD8+ T cells) and its phenotypic markers of effector ‘stemness’ (TCF-1 expression within that population), measured at an early time point after the start of rebound. This mechanistic link explains why Peluso, Sandel and Deitchman et al. observed that higher TCF-1+ fractions in cycling CD8+ T cells were associated with lower viral set points [4], e.g., individuals with a larger fraction of TCF-1+ cells within their “responding” Ki-67+ non-naïve CD8+ T cells at the post R1 time point have a greater sensitivity for effector expansion, leading to more effective viral containment. This result supports and extends our previous study, which showed that model-inferred effector expansion sensitivity predicts PTC and NC status in an ATI non-intervention setting [38], by tying that sensitivity to a measurable immunological marker. Together, they suggest that the predictive role of early responding CD8+ T cells, and of the TCF-1+ fraction within it, may generalize to other settings.

Our modeling results consistently suggest an influential role of early immune activation in determining rebound dynamics. The key immune parameter *K_B_*, related to T cell affinity for antigen [44], determines how soon and how fast effector cells expand upon encountering antigen (Fig. 4a). The maximum effector expansion rate occurs when *I* ≫ *K_B_*, as the expansion term becomes *bE*. Conversely, when *I* ≪ *K_B_*, the expansion rate is 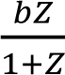. In NCs, *K_B_* is ∼ 6000 cells/mL, while for PICs, *K_B_* is ∼ 300 cells/mL (Table S2). Thus, when rebound is first starting and the number of infected cells is 100/mL, the effector expansion rate for NCs is ∼0.05/day, while for PICs it is 0.75/day, i.e. 15-fold higher. Put another way, the effector cells in NCs double every 14 days, while in the PICs they double every 0.9 days. By the time *I* = 1000 cells/mL, the effector expansion rate for NCs is 0.43/day, while for PICs it is 2.3/day, about 5 times higher, and is approaching the maximum expansion rate in the model of 3/day. This example highlights the importance of the very early post-ATI immune response: a faster ramp-up of effector activity allows the effector cell population to rapidly reach a level that can control the rebounding virus (Fig. 4a), which leads to a low steady-state level of virus and infected cells (Figs. 4b and 4c). Participants with a more sensitive CD8+ response (lower *K_B_*) effectively steered the infection toward control, whereas those with slower effector expansion (high *K_B_*) led to higher set points (Fig. 4c).

What a TCF-1-enriched responding pool contributes is then a population of T cells with a lower antigen threshold for response– consistent with the role of TCF-1+ progenitor cells in mounting secondary responses [75]. In addition, TCF-1+ cells are known to occupy a less exhausted state with reduced inhibitory signaling. While stem-like progenitor TCF-1+ CD8 T cells express PD-1, terminally exhausted TOX+ populations typically exhibit higher expression of PD-1 and additional inhibitory receptors associated with diminished TCF responsiveness [76–78]. Because PD-1 attenuates antigen receptor signaling and raises the threshold for productive activation [79,80], greater checkpoint signaling could effectively increase the antigen burden required to trigger robust proliferation. In this framework, individuals with a larger TCF-1+ population may exhibit greater expansion sensitivity because their responding cells require less antigenic stimulation to initiate expansion, consistent with the interpretation of a lower *K_B_* value.

In our modeling framework, we demonstrated mathematically that *K_B_* is linearly related to the eventual viral load set point (Equation S1, Fig. S1). This simple analytical relationship, *V*_set point_ ≈ *αK_B_*, provides a direct, testable quantitative link between a single immune parameter and clinical outcome after ATI, and a causal basis for the empirical correlation between the early responding CD8+ T cell population, with respect to both its size (%Ki-67+) and its %TCF-1+ fraction, and the viral load set point. In most participants, viral loads after the start of rebound remained highly dynamic and had not settled on a true set point during the observation period. Because Peluso, Sandel and Deitchman et al. [4] estimated the set point by averaging all data starting 2 weeks after the rebound peak, their estimate incorporates much of this variable early phase. Instead, by using the average over the last 2 weeks of observation, we found TCF-1 to be an even stronger predictor of viral control than originally reported [4]. Rutishauser et al. [75] also observed that higher %TCF-1 in HIV-specific CD8 T cells determined by MHC class I multimer staining correlates with lower VL in PWH not on ART (r ≈ –0.66). Our results extend these observations by showing that when the viral set point is defined based on endpoint values rather than the full post-peak dynamics, TCF-1’s correlation with long-term viral control is remarkably high (r ≈ –0.9, Fig. 5b). This suggests that TCF-1 is not just a correlative marker but tracks the immune-mediated control of the virus. In short, our model suggests that %TCF-1 of cycling non-naïve CD8+ T cells at early times after rebound may effectively predict the eventual set point (Fig. 5b), reinforcing its value as a surrogate indicator of post-intervention control.

Interestingly, varying the initial latent reservoir size (*L*_0_) among individuals did not significantly affect rebound dynamics in our model fits to the data. Further, the estimated value of *L*_0_ showed no between-person variation (Table S3) and had negligible impact on model outcomes (not shown). This may partly reflect the inclusion of the time-to-rebound parameter *t_a_*, which absorbs stochastic variation in the rebound event. Yet even in model variations without *t_a_*, *L*_0_ estimates remain tightly clustered, suggesting that for this cohort, the size of the inducible reservoir was not the primary driver of the post-rebound set point. This finding aligns with Peluso, Sandel and Deitchman et al.’s observations that the interventions (vaccine + bNAbs + TLR9 agonist) did not measurably reduce the reservoir, although in general all these individuals had very small reservoirs. In addition, there were no significant changes in HIV CA-DNA/RNA and that pre-ATI reservoir metrics did not correlate with rebound metrics [4]. It is also consistent with recent studies indicating that even when early ART limits reservoir size, durable control is more dependent on immune factors than on the reservoir size alone [5]. While a smaller reservoir may delay the initial time to rebound [8,9,31,81–83], our results argue that immune-mediated containment (effector cell expansion sensitivity) is the critical factor in sustaining low viremia once rebound has occurred.

The UCSF-amfAR combination immunotherapy trial did not include a placebo control arm, leaving some uncertainty as to how much the immunotherapy regimen itself contributed to the high rate of post-intervention control. However, without therapy, PTC is rare (1-6% depending on the time of ART initiation [84]) and thus one might assume the typical effector cell expansion sensitivity is too low for most individuals to control rebound and that the combination immunotherapy greatly improved the CD8+ T cell expansion potential, tipping the balance in favor of viral containment. This is consistent with the idea that the intervention “primed” the immune system in a way that standard ART alone does not. Even though Peluso, Sandel and Deitchman et al. [4] did not observe a significant vaccine-induced T cell boost or clear bNAb-mediated vaccinal effect immediately prior to rebound, the robust post-intervention control in 70% of participants suggests a more subtle immunological conditioning took place [85]. Our results hint that the treatment may have increased the pool of responsive, stem-like effector cells, which only manifest their enhanced expansion sensitivity upon viral re-exposure at ATI. For instance, without viral re-exposure post-ATI, the stem-like effector cells might stay in lymph nodes or tissues as central or tissue memory cells and thus not observable by blood sampling. This interpretation is supported by the outcome of a recent immunotherapy study in SIV-infected macaques where dual IL-10/PD-1 blockade led to an unprecedented 90% of treated animals controlling rebound, concomitant with elevated TCF-1+ memory CD8+ T cells and heightened proliferative capacity [12]. Kiani et al. [6] likewise argue that PICs already had stronger proliferative (stem-like) CD8+ features before intervention, and these features increased after administration of bNAbs [86]. These cases illustrate that therapies that enhance the sensitivity of CD8+ T cells to expand favorably upon antigen re-exposure can substantially improve remission rates. They also point to the time after ATI but before rebound as a distinct window for intervention.

There are several limitations to our study. First, because the trial lacked a control arm and included no cisgender women, we could not establish the treatment effect or determine whether the findings extend to that population. The small sample size (n = 9, PICs = 6, NCs = 3) limits the ability to obtain statistically significant comparisons between PICs and NCs in some cases. However, this is alleviated by using a mechanistic modeling framework where we compensate for the lack of statistical power with a clear mechanistic explanation. Another limitation is that our model inferences are based on viral load and CD8+ T cell data, and explicit treatment effects are not modeled. Including treatment effects is a non-trivial task since we would need to know precisely how each therapy changed model parameters for each individual. We also ignored key features of the immune system such as explicit T cell exhaustion (although it is worth noting that TCF-1+ progenitor CD8+ T cells are often the subset that avoids terminal exhaustion [87]). Another limitation is that we map total CD8+ T cells to the model effector compartment through a constant scaling parameter, which does not resolve time-varying heterogeneity in the effector fraction of circulating CD8+ T cells. Importantly, *E*(*t*) is an effective effector cell compartment rather than the HIV-specific CD8+ T cell subset. The model does not track HIV-specific cells and makes no assumption about whether their fraction of total CD8+ T cells changes during rebound.

Consistent with this, we do not know the antigen specificity of the Ki-67+ non-naïve CD8+ T cells. Finally, the estimated death rate of the cells in the effector compartment is slower than expected for a pure population of short-lived terminal effector cells (*d_E_* ≈ 0.05 per day, corresponding to a half-life of ∼2 weeks). This likely reflects that our effector compartment *E*(*t*) is an effective compartment aggregating rapidly turning-over effector cells with longer-lived progenitor/central-memory subsets (including those with TCF-1 expression) and trafficking effects. With detailed longitudinal phenotypic data, an extended model that separates effector and memory/progenitor states could provide more direct estimates of the turnover rate (*d_E_*) and even the expansion parameters (*b* and *K_B_*). Nevertheless, by approaching the problem in a more coarse-grained way, we were able to demonstrate the importance of effector expansion sensitivity to achieve post-treatment control.

In summary, our mechanistic modeling provides a clear explanation for why the responding CD8+ T cell compartment measured near the start of rebound (both the fraction of non-naïve CD8+ T cells in cycle and the TCF-1+ fraction within them) can serve as a marker of post-intervention control. It shows that both quantities track the effector cell expansion sensitivity, which is in turn inversely proportional to the eventual viral load set point, effectively translating an immunological measurement into a predictive outcome metric. This insight resolves a key uncertainty from prior clinical studies by attributing viral remission to a specific immune capability – the ability of CD8+ T cells to undergo rapid, robust expansion. Altogether, our findings emphasize effector cell expansion sensitivity, associated with the stem-like (TCF-1+) state of the responding CD8+ T cell population, as a critical determinant of HIV remission. This mechanistic understanding elevates TCF-1 and Ki-67 from correlative biomarkers to potential therapeutic targets. More broadly, our work exemplifies how simple, yet principled models can interpret complex clinical data in a biologically consistent manner, guiding the design of future cure strategies.

## Acknowledgements

This work was done under the auspices of the US Department of Energy under contract 89233218CNA000001 and supported by National Institutes of Health grants P01AI169615 and R01OD011095 (A.S.P.), R01AI152703 and U54 HL143541 (R.K.), and Martin Delaney Collaboratory UM1-AI164561 (R.M.R.). M.J.P. is supported by K23AI157875. T.P. and J.K. were partially supported by Director’s postdoctoral fellowship at Los Alamos National Laboratory (20220791PRD2 to T.K. and 20230853PRD2 to J.K.). N.P. is supported by the U.S. Department of Energy, Office of Science, Office of Advanced Scientific Computing Research, Department of Energy Computational Science Graduate Fellowship under Award Number DE-SC0022158. We also received support from The Foundation for AIDS Research (109301-59-RGRL to S.G.D.); NIH UM1AI164560 (DARE, S.G.D.), K23AI157875 (M.J.P.), R01AI170239 (R. L.R.), K23AI162249 (A.N.D.), and P30AI027763 (UCSF/Bay Area CFAR).

## Disclaimer

This report was prepared as an account of work sponsored by an agency of the United States Government. Neither the United States Government nor any agency thereof, nor any of their employees, makes any warranty, express or implied, or assumes any legal liability or responsibility for the accuracy, completeness, or usefulness of any information, apparatus, product, or process disclosed, or represents that its use would not infringe privately owned rights. Reference herein to any specific commercial product, process, or service by trade name, trademark, manufacturer, or otherwise does not necessarily constitute or imply its endorsement, recommendation, or favoring by the United States Government or any agency thereof. The views and opinions of authors expressed herein do not necessarily state or reflect those of the United States Government or any agency thereof.

## Supplementary Materials

### S1 Text. Comparison of several variations of the Conway-Perelson model

To add a non-cytolytic effect to the main model, we make the modification

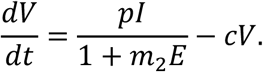

Here, *m*_2_ represents the magnitude of the effect of effector cells on reducing the rate of viral production. The model fit comparison in Table S1 shows that the main model without a noncytolytic effect is the preferred model.

**Table S1.** Model comparison.

| Model | -2LL | BICc |
| --- | --- | --- |
| *Conway – Perelson model* | 78.02 | 180.93 |
| *Main model* | **73.16** | **159.55** |
| *Main model with non-cytolytic effector function* | 80.15 | 174.81 |

### S2 Text. Best fit parameters for the main model

The best fit population and individual parameters are given in Tables S2 and S3, respectively.

**Table S2.** Best fit (population) parameters for the main model. Individual parameters are presented in Table S3. NaN indicates Monolix could not estimate the value.

| Parameter | Population estimate | 95% CI | Individual estimate (min – max) |
| --- | --- | --- | --- |
| $L_0$ (cells/mL) | 10 | NaN | [10, 10] |
| $-\log_{10} \beta$ (mL viral copies<sup>-1</sup> day<sup>-1</sup>) | 8.5 | [8.42, 8.58] | [8.50, 8.50] |
| $\lambda_E$ (cells mL<sup>-1</sup> day<sup>-1</sup>) | 1.34 | [0.76, 2.25] | [0.52, 2.04] |
| $\log_{10} K_B$ (NC) | 3.77 | [2.98, 4.55] | [3.65, 3.75] |
| $\log_{10} K_B$ (PIC) | 2.46 | [1.99, 2.92] | [1.77, 3.25] |
| $d_E$ (day<sup>-1</sup>) | 0.049 | [0.028, 0.086] | [0.018, 0.084] |
| $m$ (mL cell<sup>-1</sup> day<sup>-1</sup>) | 0.023 | NaN | [0.022, 0.025] |
| $\log_{10} p$ (day<sup>-1</sup>) | 4.12 | [4.04, 4.2] | [4.10, 4.14] |
| $t_a$ (days) | 87.54 | [59.38, 129.04] | [28.23, 170.36] |
| $f_E$ | 0.044 | NaN | [0.039, 0.050] |

**Table S3.** Best fit (individual) parameters for the main model. Population parameters and parameter units are presented in Table S2.

| *PID* | $L_0$ | $-\log_{10} \beta$ | $\lambda_E$ | $\log_{10} K_B$ | $d_E$ | $m$ | $\log_{10} p$ | $t_a$ | $f_E$ |
| --- | --- | --- | --- | --- | --- | --- | --- | --- | --- |
| 10227 | 10 | 8.50 | 1.23 | 1.77 | 0.045 | 0.022 | 4.12 | 136.89 | 0.041 |
| 71190 | 10 | 8.50 | 0.93 | 2.15 | 0.042 | 0.024 | 4.12 | 43.71 | 0.047 |
| 47019 | 10 | 8.50 | 1.23 | 2.28 | 0.047 | 0.025 | 4.11 | 96.80 | 0.050 |
| 72210 | 10 | 8.50 | 1.71 | 2.35 | 0.084 | 0.024 | 4.1 | 160.18 | 0.047 |
| 48758 | 10 | 8.50 | 0.52 | 2.87 | 0.018 | 0.023 | 4.12 | 170.36 | 0.046 |
| 30988 | 10 | 8.50 | 1.66 | 3.25 | 0.054 | 0.024 | 4.11 | 69.32 | 0.045 |
| 42894 | 10 | 8.50 | 1.52 | 3.65 | 0.043 | 0.022 | 4.14 | 141.16 | 0.039 |
| 31092 | 10 | 8.50 | 1.78 | 3.67 | 0.061 | 0.023 | 4.13 | 28.23 | 0.043 |
| 35933 | 10 | 8.50 | 2.04 | 3.75 | 0.084 | 0.023 | 4.14 | 74.43 | 0.041 |

### S3 Text. Robustness of correlations across perturbations to the model fitting procedure

During the model testing phase, we examined various perturbations to the model fitting. While different assumptions affect the goodness of fit (and/or BICc score), the correlation between 1/*K_B_* and markers of immune stemness at the time of rebound is robust (Tables S4-5).

**Table S4.** Main model fitting with varying the number of fitting parameters. (r) is Pearson’s correlation coefficient. Main model refers to the main text model with a covariate on *K_B_*. The parameters in the parentheses are fitted without random effects. First, we tested removing the random effects on each parameter. We selected the best variation, based on −2LL and the correlation strength. Then, we repeat the process by removing the random effects for a second parameter from the best variation. After two rounds (two parameters without random effects), removing the random effects from a third parameter does not improve the BICc by more than 5 points.

| Model | -2LL | BICc | (r) $1/K_B$ : Ki67 | $R^2$ | $p$-value | (r) $1/K_B$ : TCF-1 | $R^2$ | $p$-value | |
| --- | --- | --- | --- | --- | --- | --- | --- | --- | --- |
| Main Model ($L_0$) | 57.80 | 144.2 | 0.80 | 0.64 | 0.017 | 0.84 | 0.71 | 0.008 | |
| Main Model ($\beta$) | 70.06 | 156.5 | 0.81 | 0.66 | 0.014 | 0.80 | 0.64 | 0.017 | |
| Main Model ($\lambda_E$) | 56.43 | 142.8 | 0.80 | 0.64 | 0.017 | 0.85 | 0.72 | 0.008 | |
| Main Model ($d_E$) | 55.84 | 142.2 | 0.78 | 0.60 | 0.024 | 0.74 | 0.55 | 0.036 | |
| Main Model ($m$) | 67.27 | 153.7 | 0.80 | 0.65 | 0.016 | 0.75 | 0.57 | 0.031 | |
| Main Model ($p$) | 70.19 | 156.6 | 0.80 | 0.64 | 0.018 | 0.75 | 0.57 | 0.031 | |
| Main Model ($t_a$) | 158.6 | 245.0 | 0.84 | 0.71 | 0.009 | 0.67 | 0.44 | 0.071 | |
| Main Model ($f_E$) | 66.14 | 152.5 | 0.77 | 0.59 | 0.026 | 0.74 | 0.55 | 0.036 | |
| Main Model ($\lambda_E L_0$) | 57.49 | 141.7 | 0.80 | 0.64 | 0.017 | 0.84 | 0.71 | 0.008 | |
| Main Model ($\lambda_E \beta$) | 55.45 | 139.7 | 0.77 | 0.60 | 0.024 | 0.74 | 0.55 | 0.036 | |
| Main Model ($\lambda_E d_E$) | 51.32 | 135.5 | 0.76 | 0.57 | 0.030 | 0.72 | 0.51 | 0.046 | |
| Main Model ($\lambda_E m$) | 76.95 | 161.2 | 0.84 | 0.71 | 0.009 | 0.65 | 0.42 | 0.084 | |
| Main Model ($\lambda_E p$) | 53.94 | 138.1 | 0.77 | 0.59 | 0.026 | 0.73 | 0.54 | 0.038 | |
| Main Model ($\lambda_E t_a$) | 134.3 | 218.4 | 0.78 | 0.60 | 0.023 | 0.75 | 0.57 | 0.031 | |
| Main Model ($\lambda_E f_E$) | 66.14 | 150.6 | 0.78 | 0.61 | 0.022 | 0.86 | 0.73 | 0.007 | |
|  |  |  | 0.79 | 0.63 | 0.019 | 0.76 | 0.59 | 0.033 | Mean |
|  |  |  | 0.78 | 0.60 | 0.017 | 0.74 | 0.55 | 0.013 | Q<sub>1</sub> |
|  |  |  | 0.80 | 0.65 | 0.024 | 0.82 | 0.68 | 0.037 | Q<sub>3</sub> |
|  |  |  | 0.80 | 0.64 | 0.018 | 0.75 | 0.57 | 0.031 | Median |

**Table S5.** Main model fitting with fixed parameters varied by factors of 2 and 5. (r) is Pearson’s correlation coefficient. Main model refers to the main text model with a covariate on *K_B_*.

| Fixed parameter | (r) 1/$K_B$ : Ki67 | R<sup>2</sup> | p-value | (r) 1/$K_B$ : TCF-1 | R<sup>2</sup> | p-value | |
| --- | --- | --- | --- | --- | --- | --- | --- |
| $a_L \times 2$ | 0.79 | 0.62 | 0.020 | 0.73 | 0.54 | 0.039 | |
| $a_L \times 5$ | 0.83 | 0.68 | 0.012 | 0.83 | 0.69 | 0.011 | |
| $a_L \div 2$ | 0.77 | 0.59 | 0.026 | 0.74 | 0.54 | 0.037 | |
| $a_L \div 5$ | 0.76 | 0.58 | 0.029 | 0.74 | 0.55 | 0.034 | |
| $b \times 2$ | 0.81 | 0.65 | 0.016 | 0.84 | 0.70 | 0.009 | |
| $b \times 5$ | 0.82 | 0.67 | 0.013 | 0.77 | 0.59 | 0.025 | |
| $b \div 2$ | 0.77 | 0.60 | 0.025 | 0.74 | 0.55 | 0.036 | |
| $b \div 5$ | 0.77 | 0.60 | 0.025 | 0.74 | 0.55 | 0.036 | |
| $c \times 2$ | 0.78 | 0.61 | 0.023 | 0.74 | 0.54 | 0.037 | |
| $c \times 5$ | 0.80 | 0.64 | 0.018 | 0.85 | 0.73 | 0.007 | |
| $c \div 2$ | 0.83 | 0.68 | 0.012 | 0.61 | 0.37 | 0.109 | |
| $c \div 5$ | 0.80 | 0.65 | 0.016 | 0.52 | 0.27 | 0.191 | |
| $d_L \times 2$ | 0.76 | 0.58 | 0.028 | 0.74 | 0.55 | 0.036 | |
| $d_L \times 5$ | 0.76 | 0.58 | 0.028 | 0.74 | 0.55 | 0.035 | |
| $d_L \div 2$ | 0.80 | 0.64 | 0.017 | 0.84 | 0.71 | 0.009 | |
| $d_L \div 5$ | 0.80 | 0.64 | 0.018 | 0.73 | 0.53 | 0.039 | |
| $d_T \times 2$ | 0.80 | 0.63 | 0.018 | 0.85 | 0.72 | 0.007 | |
| $d_T \times 5$ | 0.81 | 0.65 | 0.016 | 0.85 | 0.72 | 0.008 | |
| $d_T \div 2$ | 0.75 | 0.56 | 0.032 | 0.49 | 0.24 | 0.222 | |
| $d_T \div 5$ | 0.72 | 0.51 | 0.045 | 0.68 | 0.46 | 0.066 | |
| $\delta \times 2$ | 0.80 | 0.64 | 0.017 | 0.85 | 0.71 | 0.008 | |
| $\delta \times 5$ | 0.85 | 0.72 | 0.008 | 0.54 | 0.29 | 0.166 | |
| $\delta \div 2$ | 0.77 | 0.60 | 0.024 | 0.74 | 0.55 | 0.035 | |
| $\delta \div 5$ | 0.76 | 0.58 | 0.027 | 0.74 | 0.55 | 0.035 | |
| $f_L \times 2$ | 0.82 | 0.68 | 0.012 | 0.77 | 0.60 | 0.025 | |
| $f_L \times 5$ | 0.76 | 0.58 | 0.029 | 0.74 | 0.55 | 0.036 | |
| $f_L \div 2$ | 0.82 | 0.67 | 0.013 | 0.77 | 0.59 | 0.025 | |
| $f_L \div 5$ | 0.82 | 0.68 | 0.012 | 0.77 | 0.59 | 0.027 | |
| $\lambda_T \times 2$ | 0.83 | 0.68 | 0.012 | 0.61 | 0.37 | 0.107 | |
| $\lambda_T \times 5$ | 0.73 | 0.54 | 0.039 | 0.67 | 0.44 | 0.071 | |
| $\lambda_T \div 2$ | 0.80 | 0.65 | 0.016 | 0.85 | 0.72 | 0.008 | |
| $\lambda_T \div 5$ | 0.81 | 0.65 | 0.016 | 0.84 | 0.71 | 0.009 | |
| $p \times 2$ | 0.81 | 0.65 | 0.015 | 0.80 | 0.65 | 0.016 | |
| $p \times 5$ | 0.86 | 0.73 | 0.007 | 0.76 | 0.58 | 0.028 | |
| $p \div 2$ | 0.82 | 0.67 | 0.013 | 0.78 | 0.60 | 0.024 | |
| $p \div 5$ | 0.76 | 0.57 | 0.030 | 0.74 | 0.55 | 0.036 | |
|  | 0.79 | 0.63 | 0.02 | 0.74 | 0.56 | 0.046 | Mean |
|  | 0.77 | 0.59 | 0.01 | 0.74 | 0.54 | 0.01 | Q<sub>1</sub> |
|  | 0.82 | 0.67 | 0.03 | 0.81 | 0.66 | 0.04 | Q<sub>3</sub> |
|  | 0.80 | 0.64 | 0.02 | 0.74 | 0.55 | 0.04 | Median |

### S4 Text. Viral set point correlation to stemness marker

Previously, we established that for the main model there is a linear relationship between *K_B_* and the viral set point [38], shown in Equation S1, and demonstrated in Fig. S1. We further observe that due to the small variations between the parameters other than *K_B_* across participants (Fig. 3a), the term in the parentheses of Equation S1 is approximately constant among participants. This gives the last approximation in equation S1, which is the basis of the correlation between *K_B_* and the viral set point.

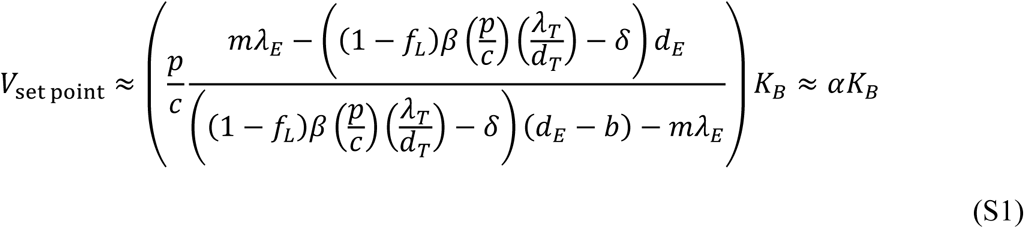

**Figure S1.**
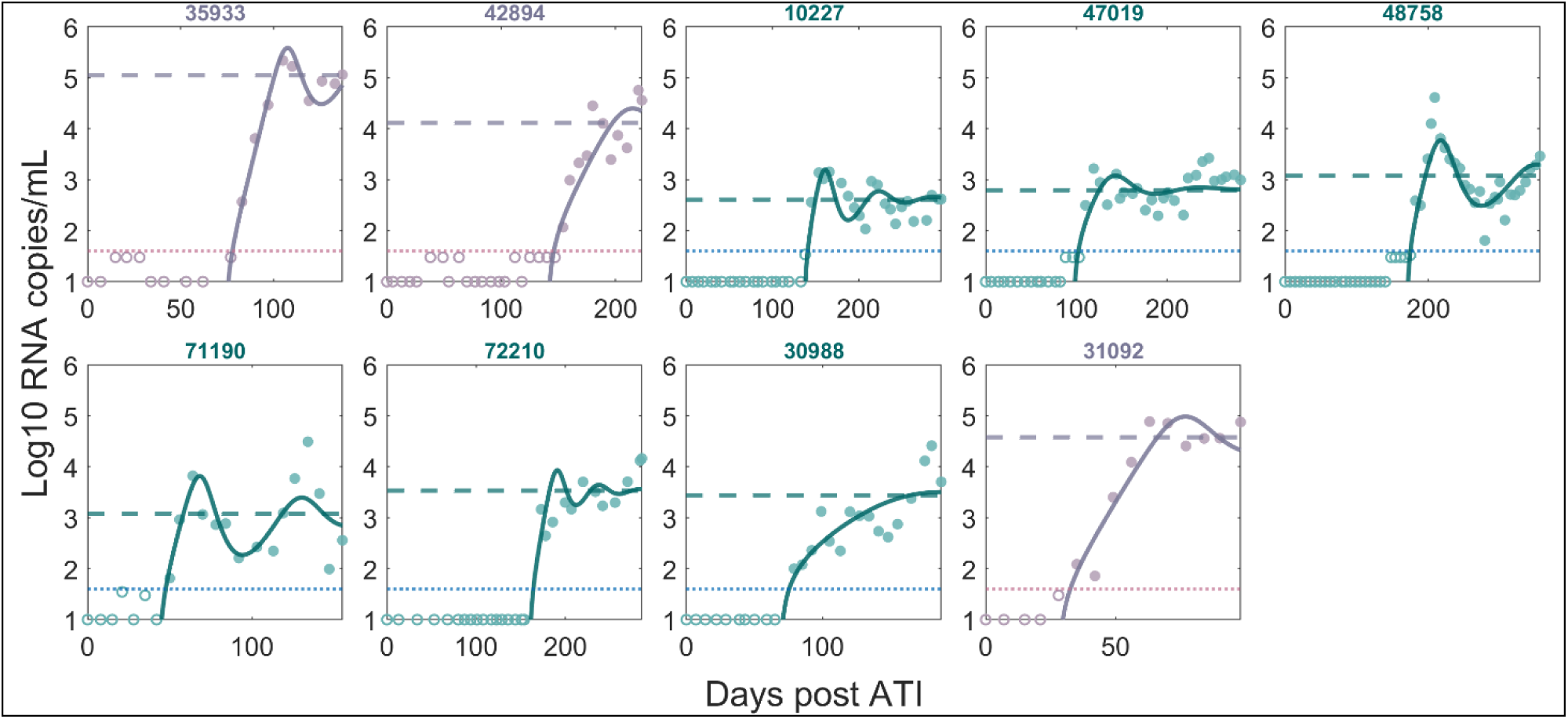
*V*_set point_ (given by Equation S1 and shown by a horizontal dashed line) approximates the viral set point well. All viral load trajectories (solid line) either approach or oscillate about *V*_set point_.

By averaging over the viral load from the last N-days (N = 7 to 84 days) of observation and using that as an estimate of the viral set point, we show that the viral set point robustly correlates with stemness markers. The correlation with %Ki67+ is strongest using the last week of viral load observations, e.g., the last 2 viral load observations, and the correlation with %TCF-1+ is strongest using the last 2 weeks of observations, e.g., the last 3 viral load observations, (Fig. S2).

**Figure S2.**
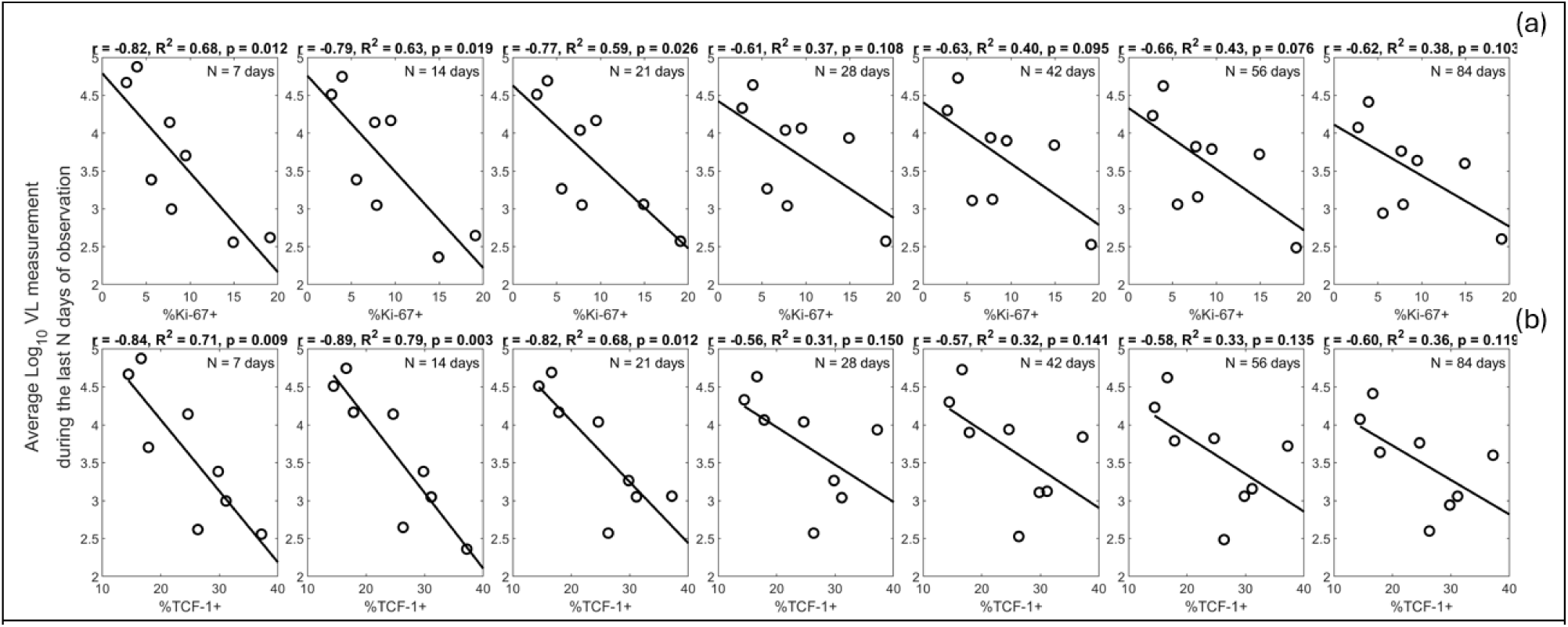
Viral set point correlations with CD8+ T cell markers. Viral set point is estimated using average viral load from the last N days of observation, where N is given at the top right of each panel. (a) Viral set point correlation with %Ki-67+ of non-naïve CD8+ T cells at post-R1. (b) Viral set point correlation with %TCF-1+ of Ki-67+ non-naïve CD8+ T cells at post-R1.

